# Detection of phylogenetic core groups in diverse microbial ecosystems

**DOI:** 10.1101/2020.01.07.896985

**Authors:** Marcos Parras-Moltó, Daniel Aguirre de Cárcer

## Abstract

The detection and subsequent analysis of phylogenetic core groups (PCGs) in a microbial ecosystem has been recently proposed as a potentially important analytical framework with which to increase our understanding of its structure and function. However, it was still unclear whether PCGs represented an infrequent phenomenon in nature. Here we provide evidence of PCGs in a large and diverse array of environments, which seems to indicate that their existence is indeed a predominant feature of microbial ecosystems. Moreover, we offer dedicated scripts to examine the presence and characteristics of PCGs in other microbial community datasets.

## Background

It is nowadays commonly believed that microbial communities assemble on the basis of function alone. This idea is supported by the predominant observation that different community compositions can translate into functionally-equivalent microbial ecosystems. In this model, multiple unrelated populations would be functionally redundant [1] in a particular microbial ecosystem type. However, this idea is somewhat challenged by the extended phenomenon of phylogenetic clustering in microbial communities; the tendency of bacteria to co-occur with phylogenetically related populations more often than expected by chance alone [2, 3].

Phylogenetic clustering in a microbial ecosystem can be studied in terms of phylogenetic core groups (PCGs), representing discrete portions of the bacterial phylogeny present in all instances of a given ecosystem type. PCGs have been detected so far in the rice rhizosphere [4] and human gut (fecal) [5] environments. The existence of a PCG in a particular microbial ecosystem type has been theorized to be linked to selection based on a combination of biotic and abiotic factors characteristic of that ecosystem, and the existence in populations belonging to that PCG of a phylogenetically-conserved set of traits allowing them to surpass such selection [4]. Thus, the study of PCGs in a given ecosystem could help understand the selection forces at play in the ecosystem, and thus illuminate overall community assembly and function.

It is still unclear whether PCGs are a predominant feature of microbial ecosystems or a rare phenomenon. Thus, to test these possibilities we evaluate here the existence of PCGs in a wide array of diverse microbial ecosystems. Also, so far PCGs had been detected in terms of 16S sequence clusters of varying depth, which represents a reasonable proxy. However, sequence clustering lacks true transitivity, which, jointly with differential initial seeding between clustering runs, may translate into slightly different clusters for the same input dataset generated by different runs or clustering algorithms. Thus, here we analyze PCGs also on the basis of nodes in a phylogenetic tree detected in all instances of the ecosystem type, an approach that also provides increased phylogenetic resolution.

## Methods

Here we analyze the existence of PCGs in nine different datasets from the literature presenting a comparatively high number of ecosystem replicates and sequencing depth (Table 1); The human microbiome is represented by datasets *FlemishGut* [6] (fecal), *TwinsUK* [7] (fecal), *Illeum* [8] (mucosa), *Rectum* [8] (mucosa), and *Vagina* [9] (mucosa). Plant-associated environments are represented by *Rice* (root samples) [10] and *Leaf* [11], animal microbiomes by *Sponge* (*Carteriospongia foliascens*) [12] and *Mice* [13], and environmental communities by *Wastewater* [14]. *Rice* was further subdivided by root environment (rhizosphere, rhizoplane, and endosphere), *Mice* by origin (wild or lab), and *Vagina* in terms of previously reported community types [9].

**Table 1.** PCGs in diverse ecosystems.

| Dataset | Samples | Depth | Frequency | Core groups per clust. threshold |
| --- | --- | --- | --- | --- |
| FlemishGut | 873 | 8,383 | 0.50±0.13 | 97<sup>2</sup>, 95<sup>1</sup>, 93<sup>1</sup>, 92<sup>1</sup>, 90<sup>1</sup>, 89<sup>3</sup>, 88<sup>3</sup>, 86<sup>3</sup>, 84<sup>1</sup>, 82<sup>1</sup>, 77<sup>1</sup> |
| TwinsUK | 2,727 | 14,082 | 0.37±0.11 | 90<sup>2</sup>, 89<sup>1</sup>, 88<sup>1</sup>, 86<sup>1</sup>, 83<sup>2</sup>, 81<sup>1</sup>, 77<sup>1</sup> |
| Illeum | 429 | 3,624 | 0.34±0.17 | 84<sup>1</sup>, 81<sup>1</sup> |
| Rectum | 304 | 3,763 | 0.42±0.18 | 86<sup>1</sup>, 79<sup>1</sup>, 75<sup>1</sup> |
| Rice | 372 | 16,884 | 0.43±0.20 | 97<sup>□</sup>, 95<sup>1</sup>, 90<sup>1</sup>, 87<sup>1</sup>, 86<sup>1</sup>, 85<sup>1</sup>, 84<sup>1</sup>, 83<sup>□</sup>, 82<sup>□</sup>, 81<sup>3</sup>, 80<sup>3</sup>, 79<sup>□</sup>, 78<sup>3</sup>, 77<sup>3</sup>, 76<sup>2</sup>, 75<sup>1</sup> |
| Rice (Rhizosphere) | 125 | 16,884 | 0.50±0.03 | 97<sup>□□</sup>, 96<sup>□</sup>, 95<sup>□</sup>, 94<sup>□</sup>, 93<sup>□</sup>, 92<sup>□</sup>, 91<sup>□</sup>, 90<sup>□</sup>, 89<sup>1□</sup>, 88<sup>1□</sup>, 87<sup>1□</sup>, 86<sup>1□</sup>, 85<sup>□</sup>, 84<sup>1□</sup>, 83<sup>12</sup>, 82<sup>□</sup>, 81<sup>□</sup>, 80<sup>□</sup>, 79<sup>13</sup>, 78<sup>□</sup>, 77<sup>□</sup>, 76<sup>□</sup>, 75<sup>1</sup> |
| Rice (Endosphere) | 133 | 17,735 | 0.70±0.15 | 97<sup>1□</sup>, 96<sup>1</sup>, 95<sup>1</sup>, 92<sup>1</sup>, 91<sup>1</sup>, 90<sup>1</sup>, 88<sup>1</sup>, 87<sup>2</sup>, 86<sup>3</sup>, 85<sup>2</sup>, 84<sup>2</sup>, 83<sup>1</sup>, 82<sup>□</sup>, 81<sup>□</sup>, 80<sup>2</sup>, 79<sup>2</sup>, 78<sup>2</sup>, 77<sup>2</sup>, 76<sup>2</sup>, 75<sup>3</sup> |
| Rice (Rhizoplane) | 114 | 17,821 | 0.54±0.09 | 97<sup>33</sup>, 96<sup>□</sup>, 95<sup>3</sup>, 94<sup>3</sup>, 93<sup>3</sup>, 92<sup>□</sup>, 91<sup>□</sup>, 90<sup>□</sup>, 89<sup>2</sup>, 88<sup>□</sup>, 87<sup>11</sup>, 86<sup>□</sup>, 85<sup>□</sup>, 84<sup>11</sup>, 83<sup>1□</sup>, 82<sup>□</sup>, 81<sup>□</sup>, 80<sup>□</sup>, 79<sup>□</sup>, 78<sup>□</sup>, 77<sup>□</sup>, 76<sup>1</sup>, 75<sup>2</sup> |
| Vagina (#1-3,5) | 286 | 693 | 0.93±0.11 | 82<sup>1</sup> |
| Vagina (#1) | 105 | 693 | 0.80±0.19 | 97<sup>1</sup> |
| Vagina (#2) | 25 | 774 | 0.79±0.17 | 97<sup>1</sup> |
| Vagina (#3) | 135 | 1,271 | 0.85±0.13 | 97<sup>1</sup> |
| Vagina (#4) | 108 | 881 | NA | NA |
| Vagina (#5) | 21 | 936 | 0.76±0.17 | 97<sup>1</sup>, 95<sup>1</sup> |
| Wastewater | 43 | 48,668 | 0.53±0.07 | 97<sup>□□</sup>, 96<sup>2</sup>, 95<sup>2</sup>, 94<sup>1</sup>, 93<sup>□</sup>, 92<sup>□</sup>, 91<sup>□</sup>, 90<sup>□</sup>, 89<sup>□</sup>, 88<sup>□</sup>, 87<sup>□</sup>, 86<sup>1□</sup>, 85<sup>□</sup>, 84<sup>□</sup>, 83<sup>11</sup>, 82<sup>3</sup>, 81<sup>□</sup>, 8<sup>3</sup>, 79<sup>3</sup>, 78<sup>□</sup>, 77<sup>3</sup>, 76<sup>□</sup>, 75<sup>□</sup> |
| Sponge | 143 | 27,921 | 0.77±0.10 | 97<sup>1□</sup>, 96<sup>1</sup>, 93<sup>1</sup>, 92<sup>3</sup>, 91<sup>1</sup>, 90<sup>3</sup>, 89<sup>1</sup>, 88<sup>□</sup>, 87<sup>□</sup>, 86<sup>□</sup>, 85<sup>1</sup>, 84<sup>□</sup>, 83<sup>2</sup>, 81<sup>3</sup>, 80<sup>3</sup>, 78<sup>2</sup>, 76<sup>1</sup> |
| Leaf | 175 | 10,322 | 0.18±0.10 | 97<sup>1</sup>, 91<sup>1</sup> |
| Mice (Total) | 230 | 10,086 | 0.52±0.08 | 95<sup>1</sup>, 92<sup>1</sup>, 91<sup>2</sup>, 91<sup>1</sup>, 89<sup>2</sup>, 88<sup>3</sup>, 87<sup>2</sup>, 86<sup>2</sup>, 85<sup>□</sup>, 84<sup>□</sup>, 83<sup>□</sup>, 82<sup>□</sup>, 81<sup>1</sup>, 80<sup>1</sup>, 79<sup>1</sup>, 78<sup>1</sup> |
| Mice (Lab) | 129 | 15,932 | 0.51±0.10 | 97<sup>21</sup>, 96<sup>3</sup>, 95<sup>2</sup>, 94<sup>2</sup>, 93<sup>2</sup>, 92<sup>□</sup>, 91<sup>1□</sup>, 90<sup>1□</sup>, 89<sup>□</sup>, 88<sup>□</sup>, 87<sup>□</sup>, 86<sup>□</sup>, 85<sup>3</sup>, 84<sup>3</sup>, 83<sup>□</sup>, 81<sup>□</sup>, 80<sup>3</sup>, 79<sup>1</sup>, 78<sup>1</sup> |
| Mice (Wild) | 101 | 10,086 | 0.57±0.07 | 97<sup>3</sup>, 96<sup>1</sup>, 93<sup>1</sup>, 92<sup>2</sup>, 90<sup>2</sup>, 89<sup>1</sup>, 88<sup>2</sup>, 87<sup>□</sup>, 86<sup>□</sup>, 85<sup>□</sup>, 84<sup>2</sup>, 83<sup>1</sup>, 82<sup>1</sup>, 81<sup>1</sup>, 79<sup>1</sup>, 78<sup>2</sup>, 77<sup>1</sup>, 75<sup>1</sup> |

**Depth**; sequences per sample. **Frequency**; average pooled abundance of members of the core OTUs across the dataset. **Core groups per clust. threshold**; Numbers represent similarity clustering thresholds (x10<sup>-2</sup>) were core OTUs were detected, and superscript values indicate the number of such OTUs observed for each threshold.

For each dataset, samples presenting very low sequence depths were removed, then all samples were subsampled to a (minimum) common depth. Finally, the normalized datasets were analyzed with *BacterialCore*.*py* (https://git.io/Je5V3). The script uses various QIIME processes [15] and *R* libraries to detect PCGs and produce associated analyses and statistics. It employs the clustering-based core detection approach previously described [5], and a new approach based on a 16S rRNA gene phylogeny. Here, the algorithm traverses the tree from leaves to root; if a leaf/node is present in a (selected) percentage of samples it is flagged as “core”, and its abundance values removed from all parental nodes before continuing, so that reported core groups are non-overlapping. Additionally, *BacterialCore*.*py* provides per core-group information, statistics, and consensus taxonomies.

## Results and discussion

The microbial ecosystems analyzed presented a considerable number of PCGs detected at different phylogenetic depths along the bacterial phylogeny (Table 1, Figure 1, Suppl. Mat. 1, Suppl. Mat. 2). The exceptions to this pattern were the mucosa environments analyzed (*Illeum, Rectum*, and *Vagina*) as well as the *Leaf* ecosystem, featuring the presence of very few PCGs. This phenomenon could be hypothesized to relate to the more homogeneous abiotic conditions of these environments translating to less diverse communities. However, the *Rice rhizoplane* and *endosphere* ecosystems, which could also be *a priori* considered as presenting more homogeneous abiotic conditions, presented a large number of PCGs. The low number of PCGs detected in the mucosal ecosystems could be related to their comparatively low sequencing depth. Nevertheless, the leaf environment presents a substantial sequencing depth, but only two PCGs.

**Figure 1.**
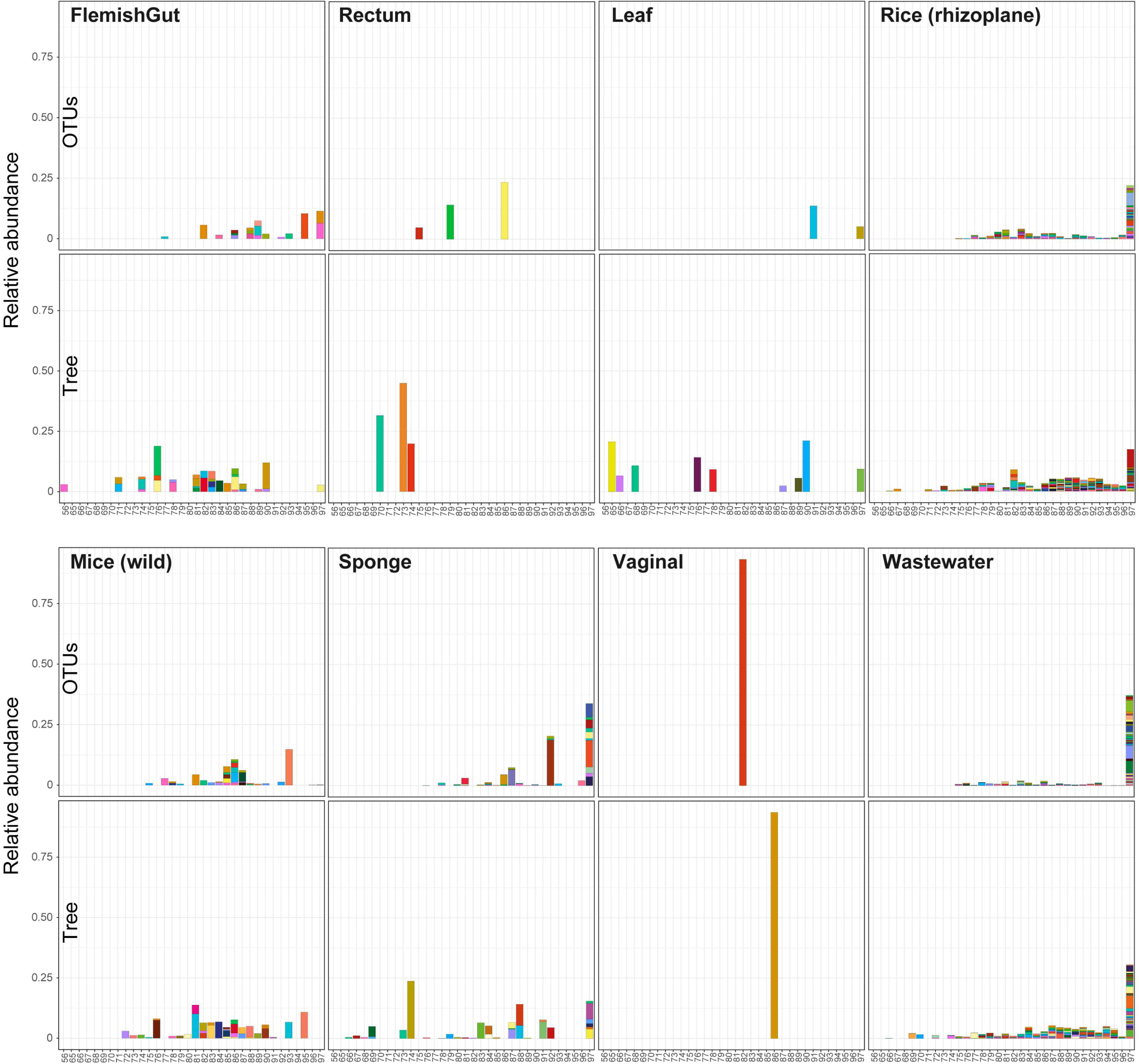
Detection of PCGs in datasets. Results for selected datasets based on the dynamic clustering of 16S rRNA gene sequences from 97% to 75% sequence identity (right to left) [OTUs] and the phylogenetic tree-based approach [Tree]. For each threshold, OTUs/nodes present in all samples (i.e. core) appear vertically stacked with individual heights representing average relative abundance of each core OTU/node in the dataset. For the tree-based approach, x-axis values represent the maximum intra-node distance, not the average.

Overall, the detected PCGs represented a preeminent fraction of the total community (Table 1), with the lowest pooled abundance values being 18.5% (*Leaf*) and 34.9% (*Illeum*), and the largest 77.6% (*Sponge*) and 93.4% (*Vagina*). In general, there was a good correspondence between the clustering and tree-based approaches (Figure 1, Supplementary Material 2), both of which produced correlated results in terms of number of PCGs and their phylogenetic depth. Commonly, results for the clustering approach represented a subset of those from the tree-based approach (Supplementary Material 1; Venn diagrams)

In this brief report we have detected PCGs in terms of 16S sequence clusters and nodes in a phylogenetic tree of different depths present in all samples from the same ecosystem type. While this is a useful heuristic, other criteria such as a Poisson distribution [16], a competitive lottery schema [17], invariance metrics [18], or the use of neutral models [19, 20], could be employed and implemented within *BacterialCore*.*py*.

## Conclusion

The use of observed phylogenetic clustering patterns of community assembly may represent an important clue to understand the assembly and function of a microbial ecosystem. Here we provide evidence of PCGs in a large and diverse array of environments, which seems to indicate that their existence is indeed a predominant feature of microbial ecosystems. Moreover, we offer dedicated scripts to examine the presence and characteristics of PCGs in other microbial community datasets.

## Availability of data

The datasets analyzed during the current study are available from their original sources. Additional result files and scripts are available from the corresponding author upon request.

## Acknowledgements

This work was funded by the Spanish Ministry of Science and Innovation grant BIO2016-80101-R.

## SUPPLEMENTARY MATERIALS

**Supplementary Material 1**. *BacterialCore*.*py* result files (intermediate clustering files have been omitted due to their large size).

**Supplementary Material 2**. Detection of PCGs in datasets. Results based on the dynamic clustering of 16S rRNA gene sequences from 97% to 75% sequence identity (right to left) [Page 1; OTUs], and the phylogenetic tree-based approach where x-axis values represent the maximum intra-node distance [Page 2; MaxS] or the average intra-node distance [Page3; MeanS]. For each threshold, OTUs/nodes present in all samples (i.e. core) appear vertically stacked with individual heights representing average relative abundance of each core OTU/node in the dataset. Results arising from both approaches are also compared [Pages 4-5].

## Notes

https://git.io/Je5V3

## REFERENCES

1. Adair KL, Douglas AE. Making a microbiome: the many determinants of host-associated microbial community composition. Curr Opin Microbiol. 2017;35:23–9.

2. Stegen JC, Lin X, Konopka AE, Fredrickson JK. Stochastic and deterministic assembly processes in subsurface microbial communities. The Isme Journal. 2012;6:1653.

3. Horner-Devine MC, Bohannan BJ. Phylogenetic clustering and overdispersion in bacterial communities. Ecology. 2006;87:S100–8.

4. Aguirre de Cárcer D. A conceptual framework for the phylogenetically constrained assembly of microbial communities. Microbiome. 2019;7:142.

5. Aguirre de Cárcer D. The human gut pan-microbiome presents a compositional core formed by discrete phylogenetic units. Scientific Reports. 2018;8:14069.

6. Falony G, Joossens M, Vieira-Silva S, Wang J, Darzi Y, Faust K et al. Population-level analysis of gut microbiome variation. Science. 2016;352:560–4.

7. Goodrich JK, Davenport ER, Beaumont M, Jackson MA, Knight R, Ober C et al. Genetic Determinants of the Gut Microbiome in UK Twins. Cell Host Microbe. 2016;19:731–43.

8. Gevers D, Kugathasan S, Denson LA, Vazquez-Baeza Y, Van Treuren W, Ren B et al. The treatment-naive microbiome in new-onset Crohn’s disease. Cell Host Microbe. 2014;15:382–92.

9. Ravel J, Gajer P, Abdo Z, Schneider GM, Koenig SSK, McCulle SL et al. Vaginal microbiome of reproductive-age women. Proceedings of the National Academy of Sciences. 2011;108:4680–7.

10. Edwards J, Johnson C, Santos-Medellín C, Lurie E, Podishetty NK, Bhatnagar S et al. Structure, variation, and assembly of the root-associated microbiomes of rice. Proceedings of the National Academy of Sciences. 2015;112:E911.

11. Wagner MR, Lundberg DS, Del Rio TG, Tringe SG, Dangl JL, Mitchell-Olds T. Host genotype and age shape the leaf and root microbiomes of a wild perennial plant. Nat Commun. 2016;7.

12. Moitinho-Silva L, Nielsen S, Amir A, Gonzalez A, Ackermann GL, Cerrano C et al. The sponge microbiome project. Gigascience. 2017;6:1–7.

13. Rosshart SP, Vassallo BG, Angeletti D, Hutchinson DS, Morgan AP, Takeda K et al. Wild Mouse Gut Microbiota Promotes Host Fitness and Improves Disease Resistance. Cell. 2017;171:1015-28.e13.

14. Saunders AM, Albertsen M, Vollertsen J, Nielsen PH. The activated sludge ecosystem contains a core community of abundant organisms. ISME J. 2016;10:11–20.

15. Caporaso JG, Kuczynski J, Stombaugh J, Bittinger K, Bushman FD, Costello EK et al. QIIME allows analysis of high-throughput community sequencing data. Nat Meth. 2010;7:335–6.

16. Gumiere T, Meyer K, Burns A, Gumiere S, Bohannan B, Andreote F: A probabilistic model to identify the core microbial community; 2018.

17. Verster AJ, Borenstein E. Competitive lottery-based assembly of selected clades in the human gut microbiome. Microbiome. 2018;6:186.

18. Bradley PH, Pollard KS. Proteobacteria explain significant functional variability in the human gut microbiome. Microbiome. 2017;5:017–0244.

19. Harris K, Parsons TL, Ijaz UZ, Lahti L, Holmes I, Quince C. Linking Statistical and Ecological Theory: Hubbell’s Unified Neutral Theory of Biodiversity as a Hierarchical Dirichlet Process. Proceedings of the IEEE. 2017;105:516–29.

20. Adair KL, Wilson M, Bost A, Douglas AE. Microbial community assembly in wild populations of the fruit fly Drosophila melanogaster. The Isme Journal. 2018;12:959–72.

