## Supplementary material 1 for "Detection of phylogenetic core groups in diverse microbial ecosystems": README_Results.txt

#### READ ME #### 

######Supplementary results for: "Detection of phylogenetic core groups in diverse microbial ecosystems" Parras-Moltó & Aguirre de Cárcer, 2019. #######


##Output files for each dataset:

OTUs (folder):
	Statistics.txt: Total number, pooled mean abundance, standard deviation, minimun and maximun of core groups for all clustering thresholds. 
	General.txt: Number of core OTUs per clustering threshold.
	filt.txt: Reads belonging to core OTUs.
	Distr.txt: Frequency mean, Frequency standard deviation, Frequency CV, ecdf-mean and ecdf-CV for each core at each clustering level. (ecdf: rank of the value when compared to all OTUs at the same clustering thresholds)
	Core_Otus_freq.txt: Sample abundance table for each core OTU at each thresholds.
	control_random_table_cores: Number of core OTUs obtained at each thresholds after randomizing the OTU abundance table in each thresholds 1000 times.
	
Arbol (folder):
	abundances.txt: Sample abundance table for each core node/leaf.
	results.txt: Core name, absolute abundances (vector), relative abundances (vector), min relative abundance, max relative abundance, average relative abundance, SD relative abundance, consensuns taxonomy, number of leaves. 
	results_stats.txt: Number of leaves, mean 16S distance between leaves, estandard deviation of 16S distances between leaves, and max 16S distance between leaves, for each node and leaf core.

Reads_cores_Tree_ALL_OTUs_reads_core_ALL_.png #Image representing the Venn diagram (shared reads belonging to core OTUs/nodes).
