## Supplementary figures and images for "Detection of phylogenetic core groups in diverse microbial ecosystems"

### Reads_cores_Tree_ALL_OTUs_reads_core_ALL_.png

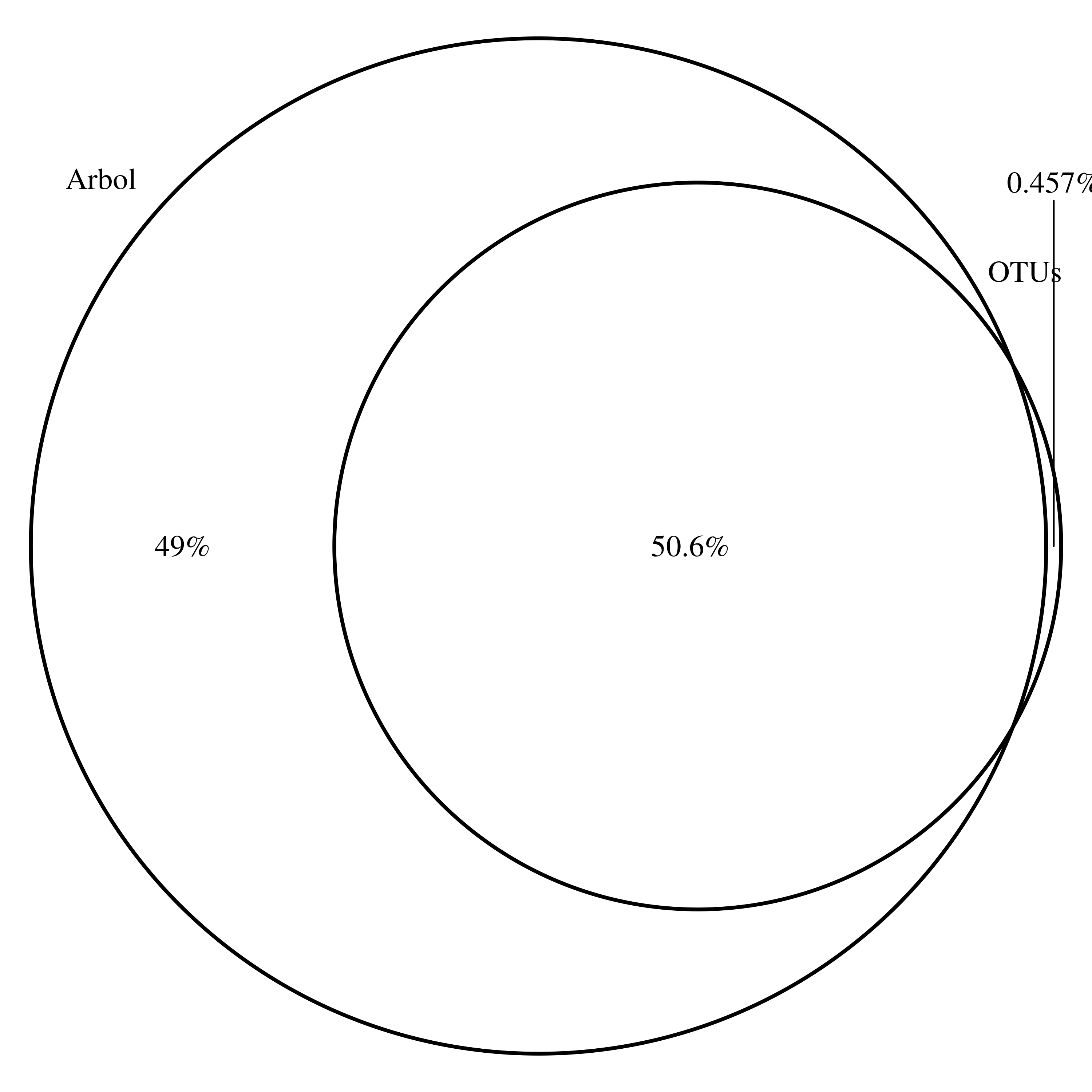

### Reads_cores_Tree_ALL_OTUs_reads_core_ALL_.png

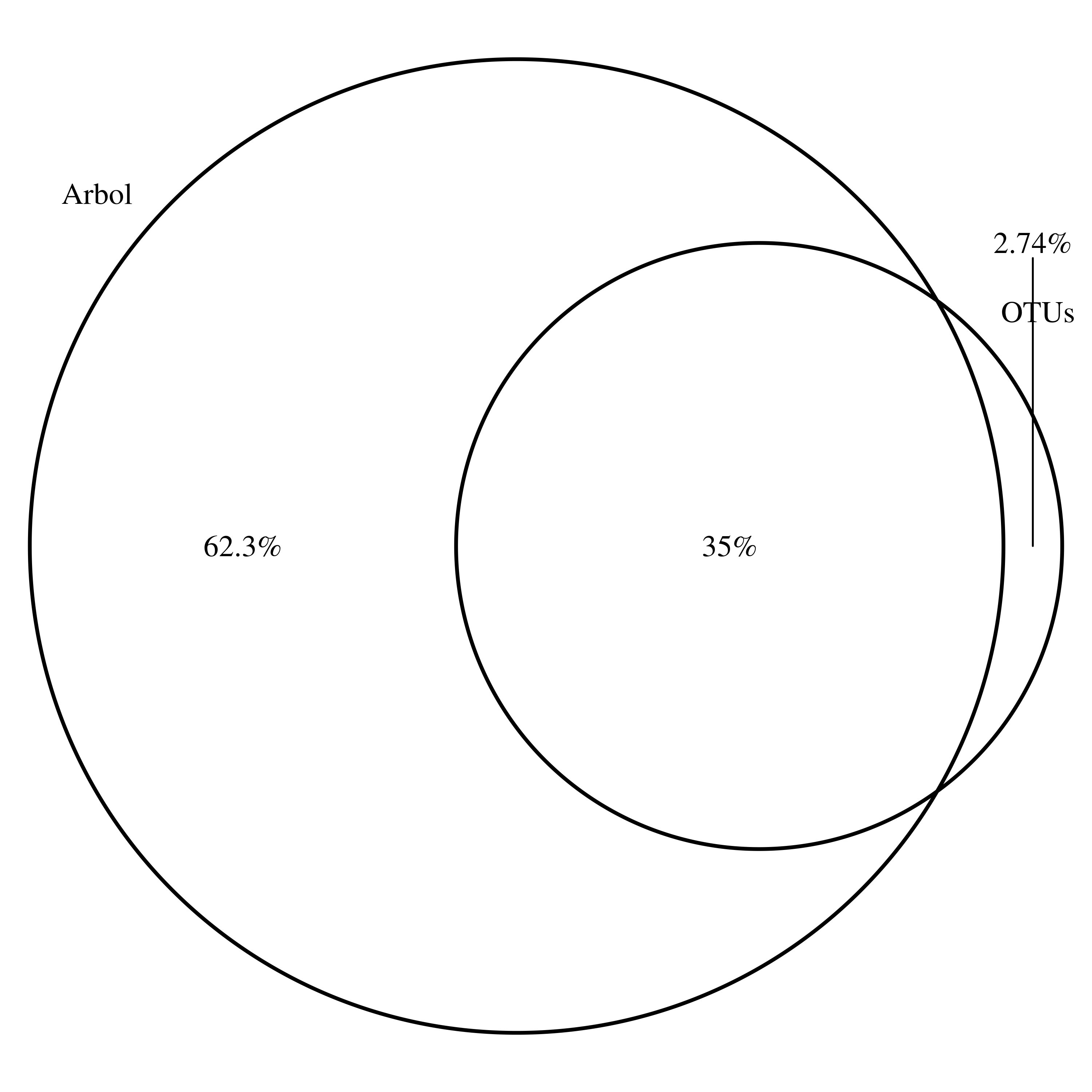

### Reads_cores_Tree_ALL_OTUs_reads_core_ALL_.png

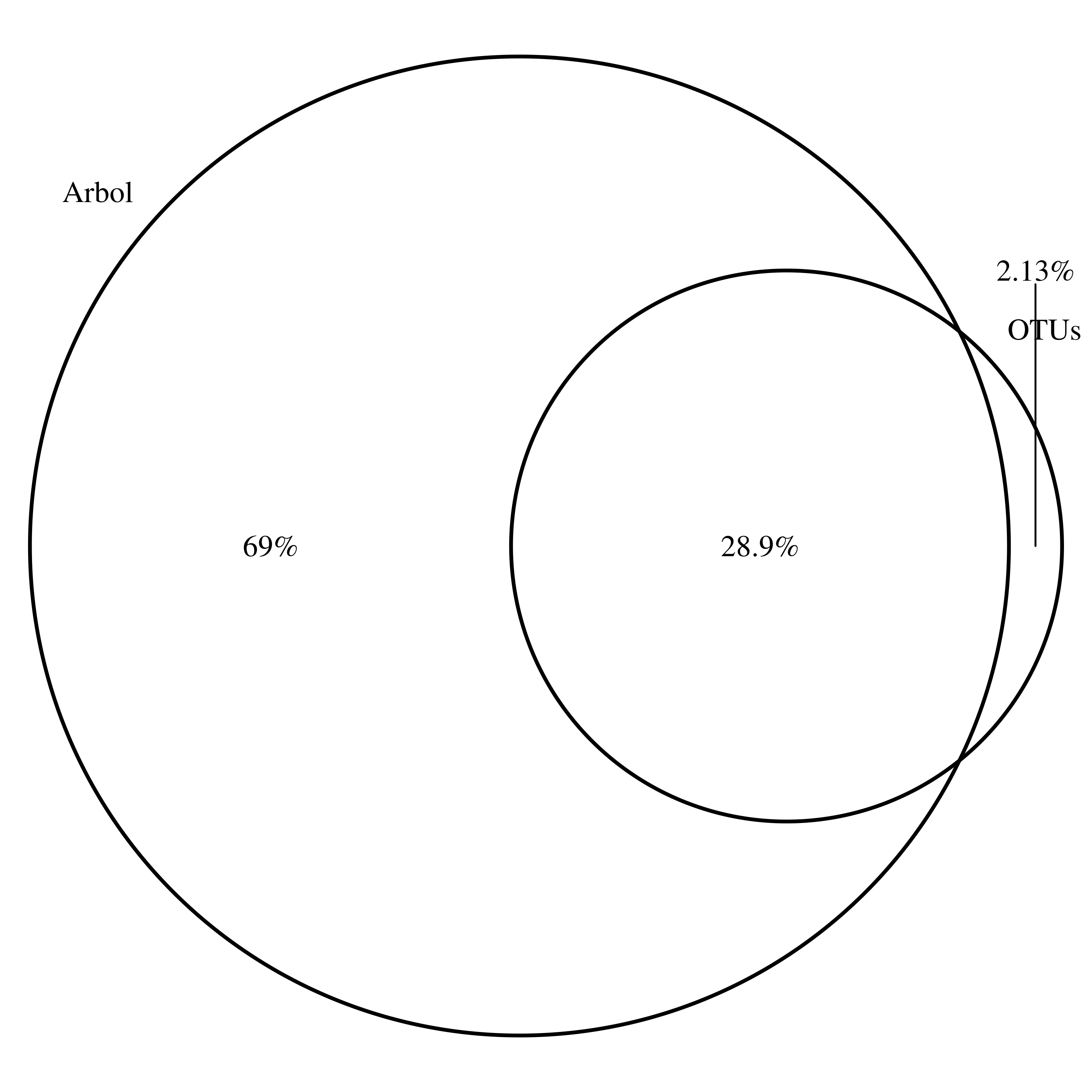

### Reads_cores_Tree_ALL_OTUs_reads_core_ALL_.png

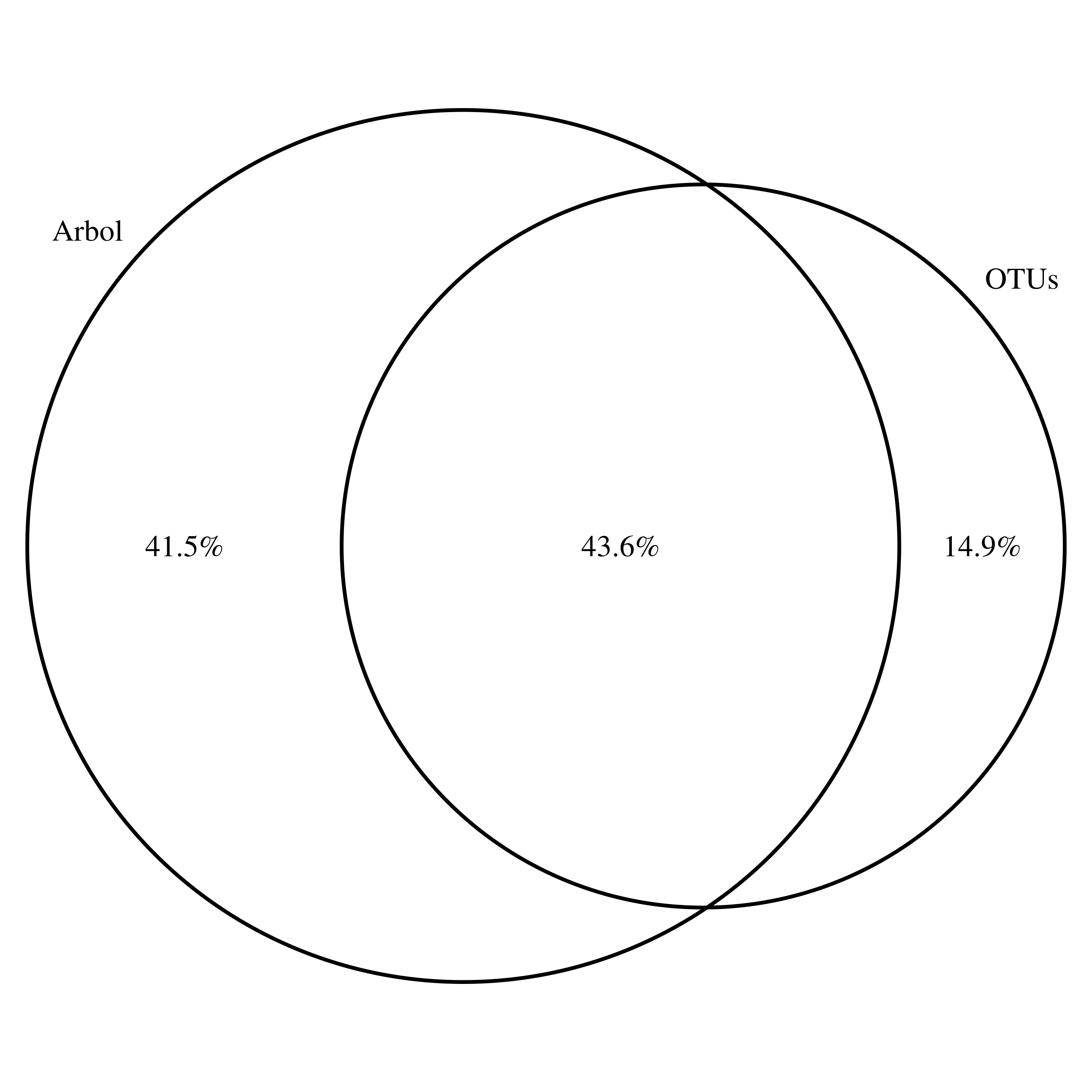

### Reads_cores_Tree_ALL_OTUs_reads_core_ALL_.png

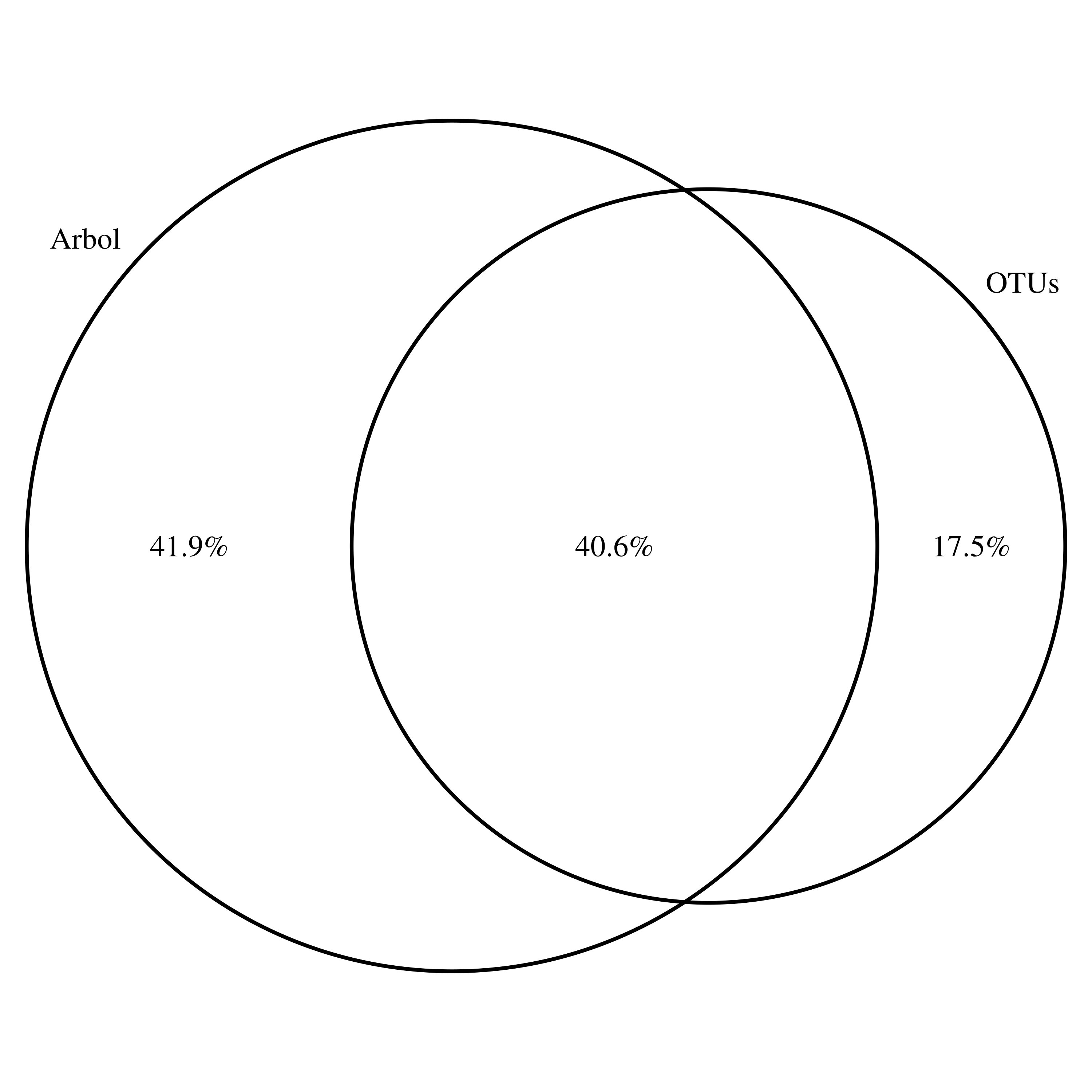

### Reads_cores_Tree_ALL_OTUs_reads_core_ALL_.png

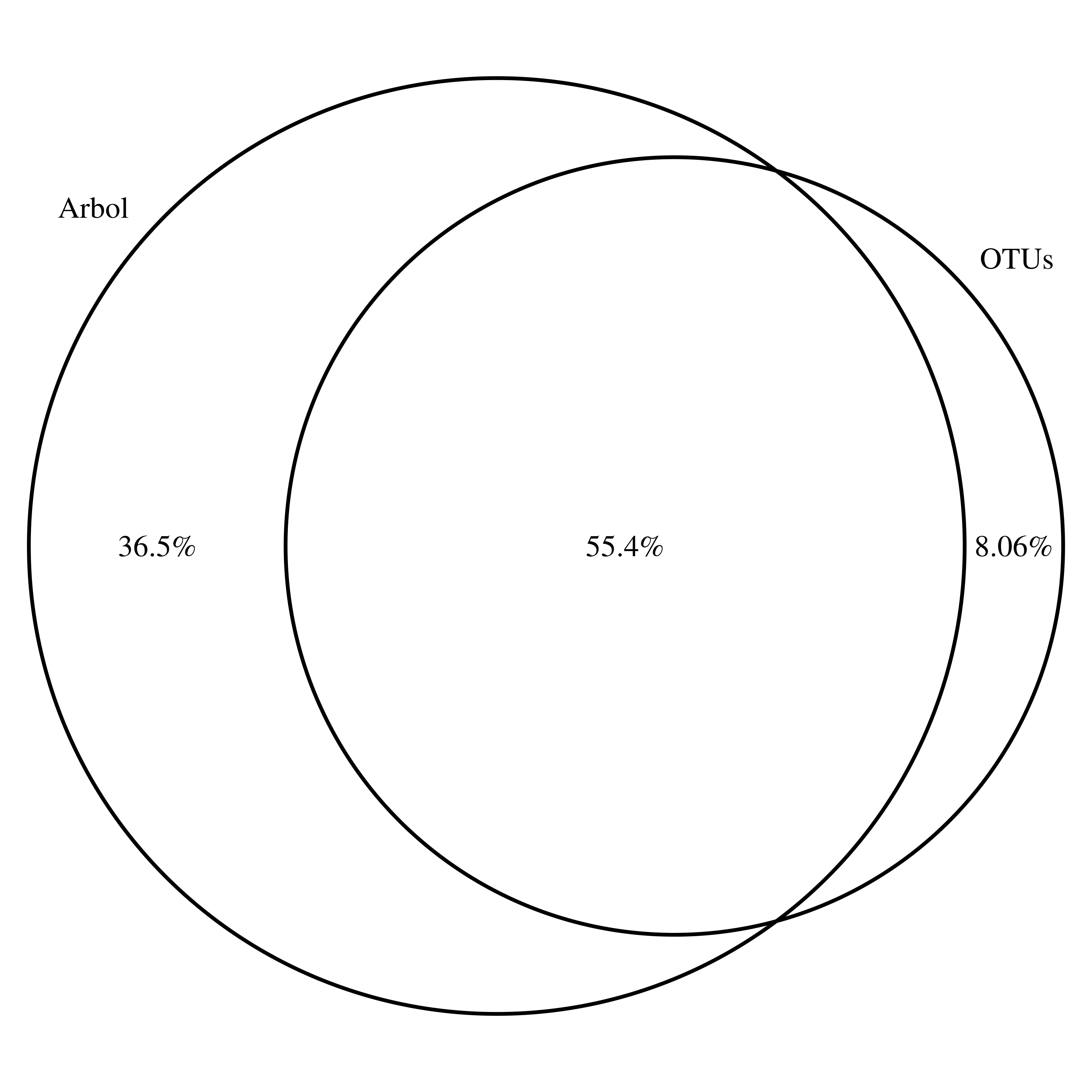

### Reads_cores_Tree_ALL_OTUs_reads_core_ALL_.png

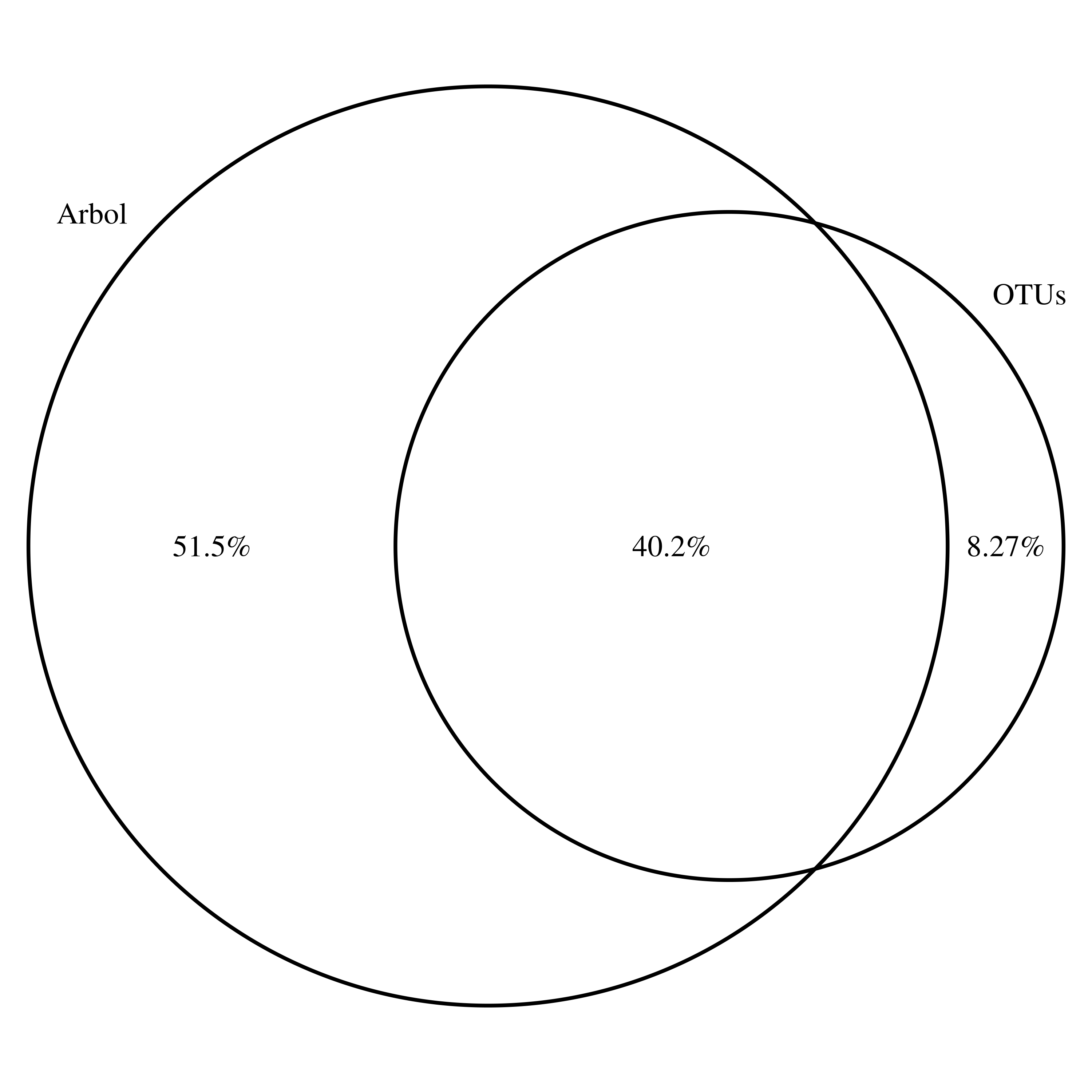

### Reads_cores_Tree_ALL_OTUs_reads_core_ALL_.png

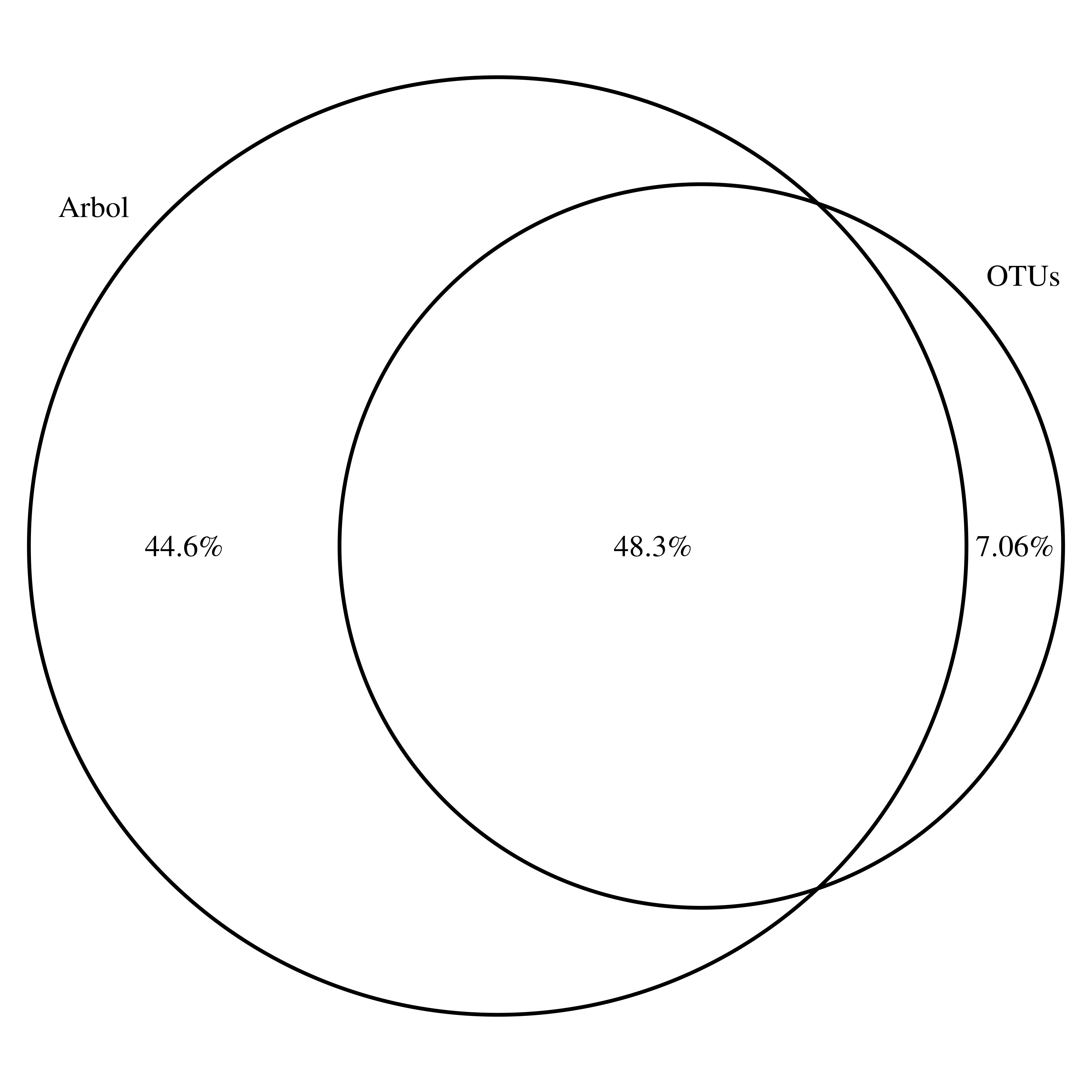

### Reads_cores_Tree_ALL_OTUs_reads_core_ALL_.png

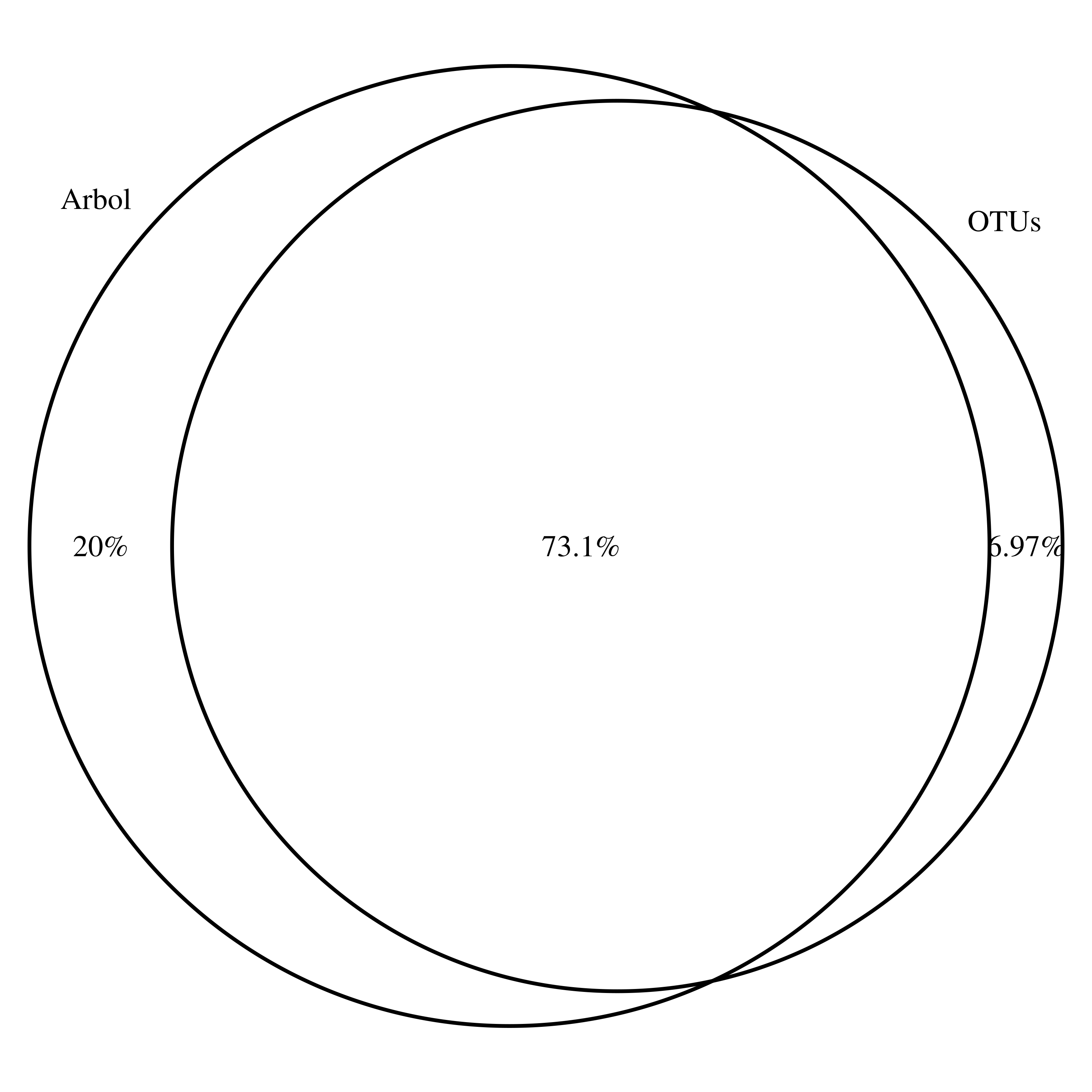

### Reads_cores_Tree_ALL_OTUs_reads_core_ALL_.png

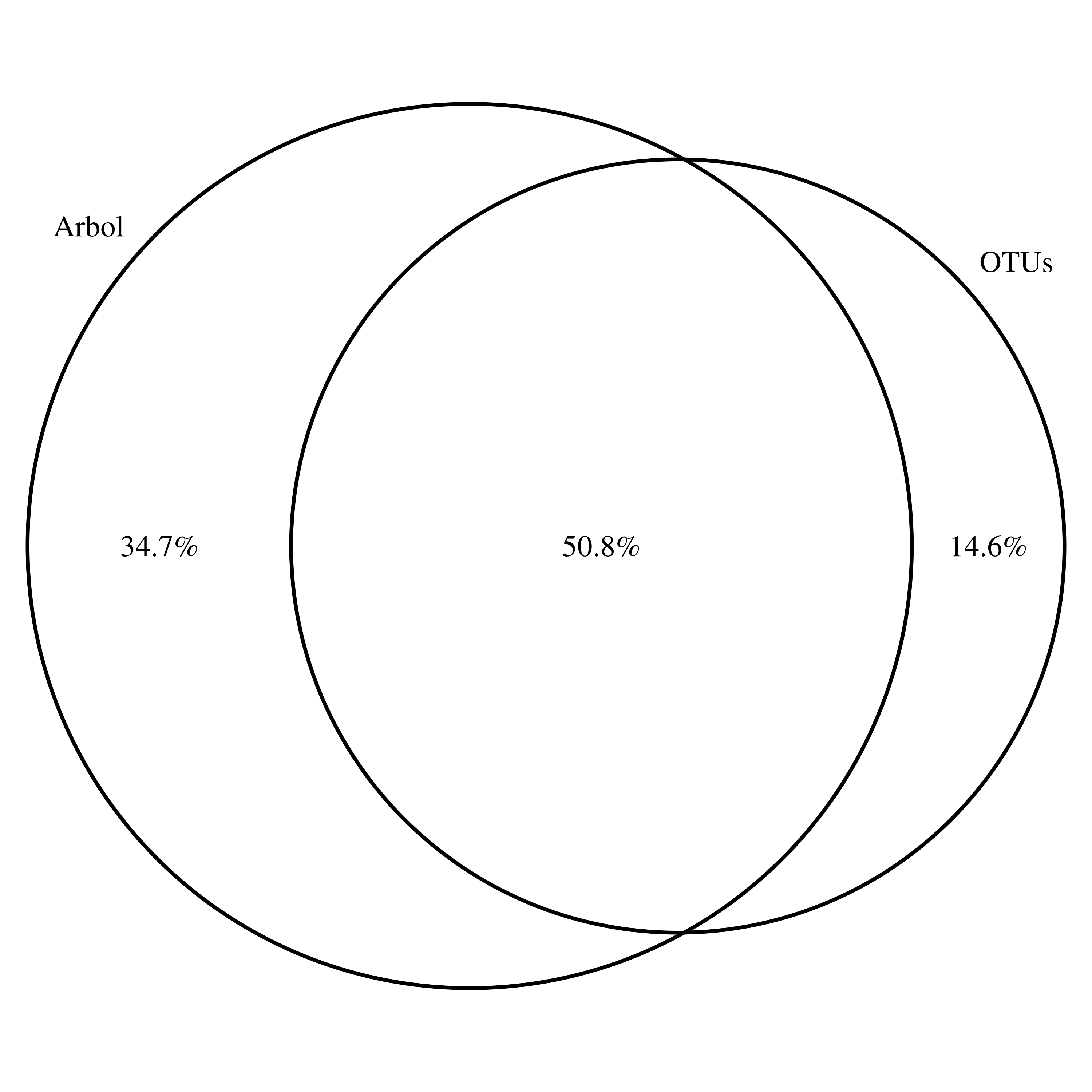

### Reads_cores_Tree_ALL_OTUs_reads_core_ALL_.png

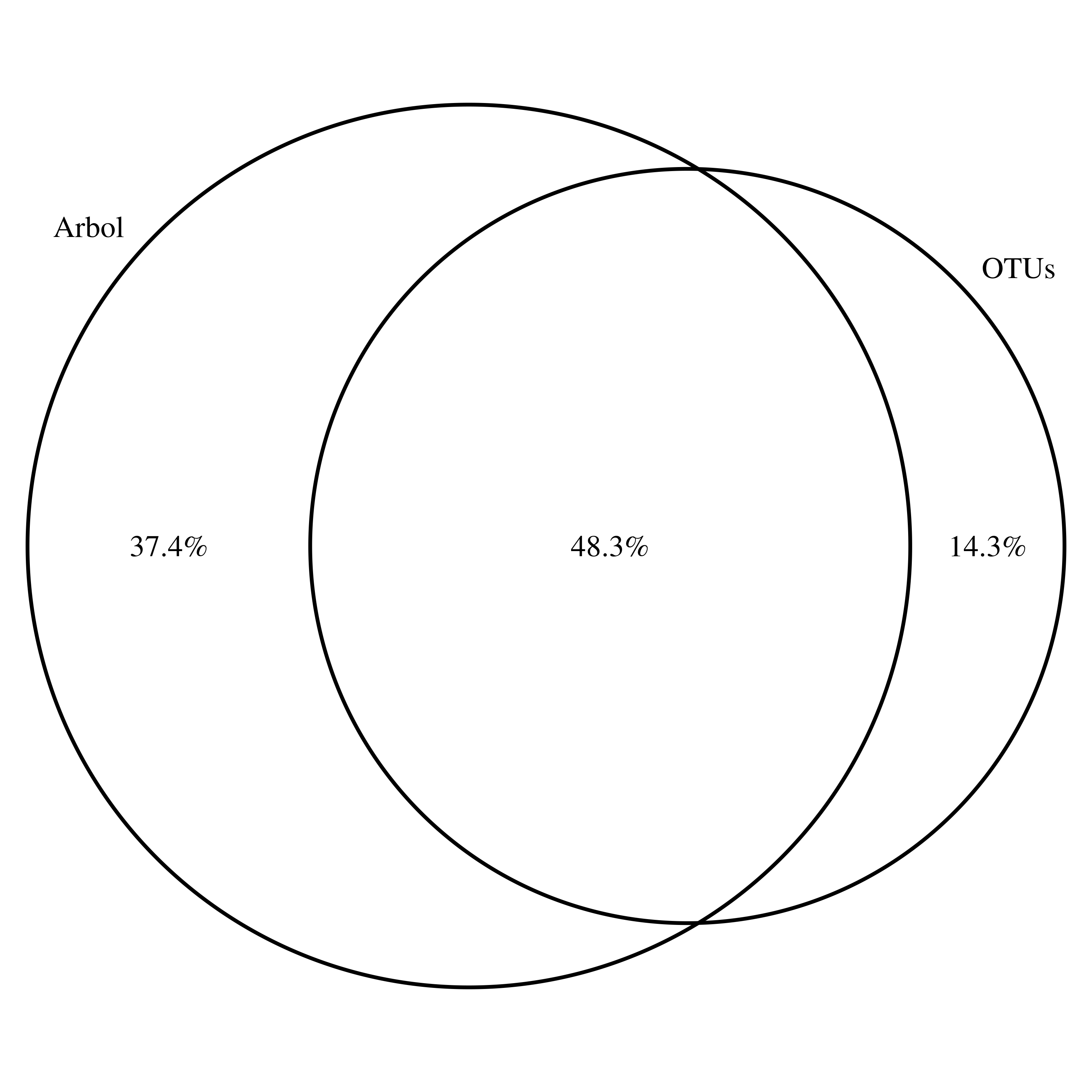

### Reads_cores_Tree_ALL_OTUs_reads_core_ALL_.png

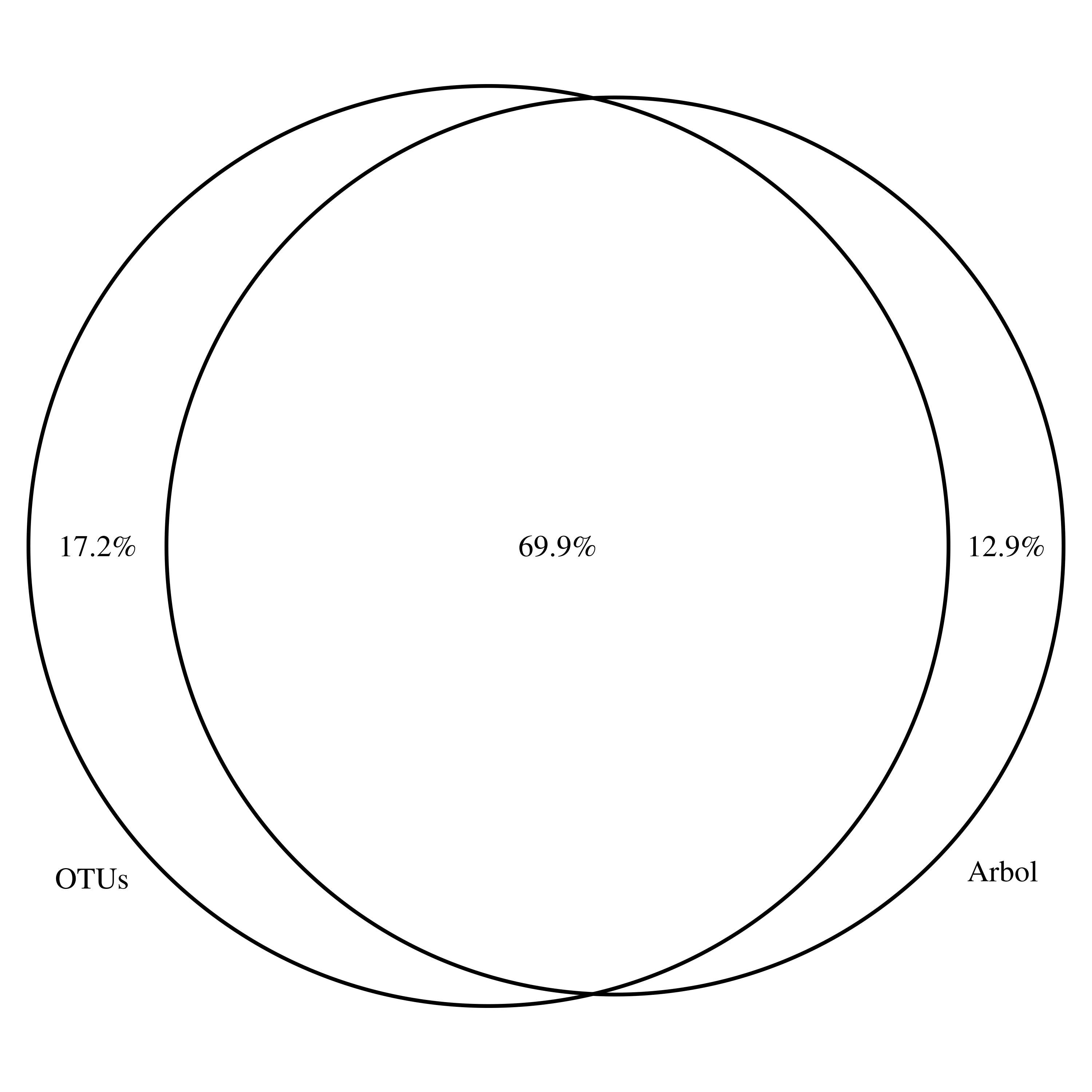

### Reads_cores_Tree_ALL_OTUs_reads_core_ALL_.png

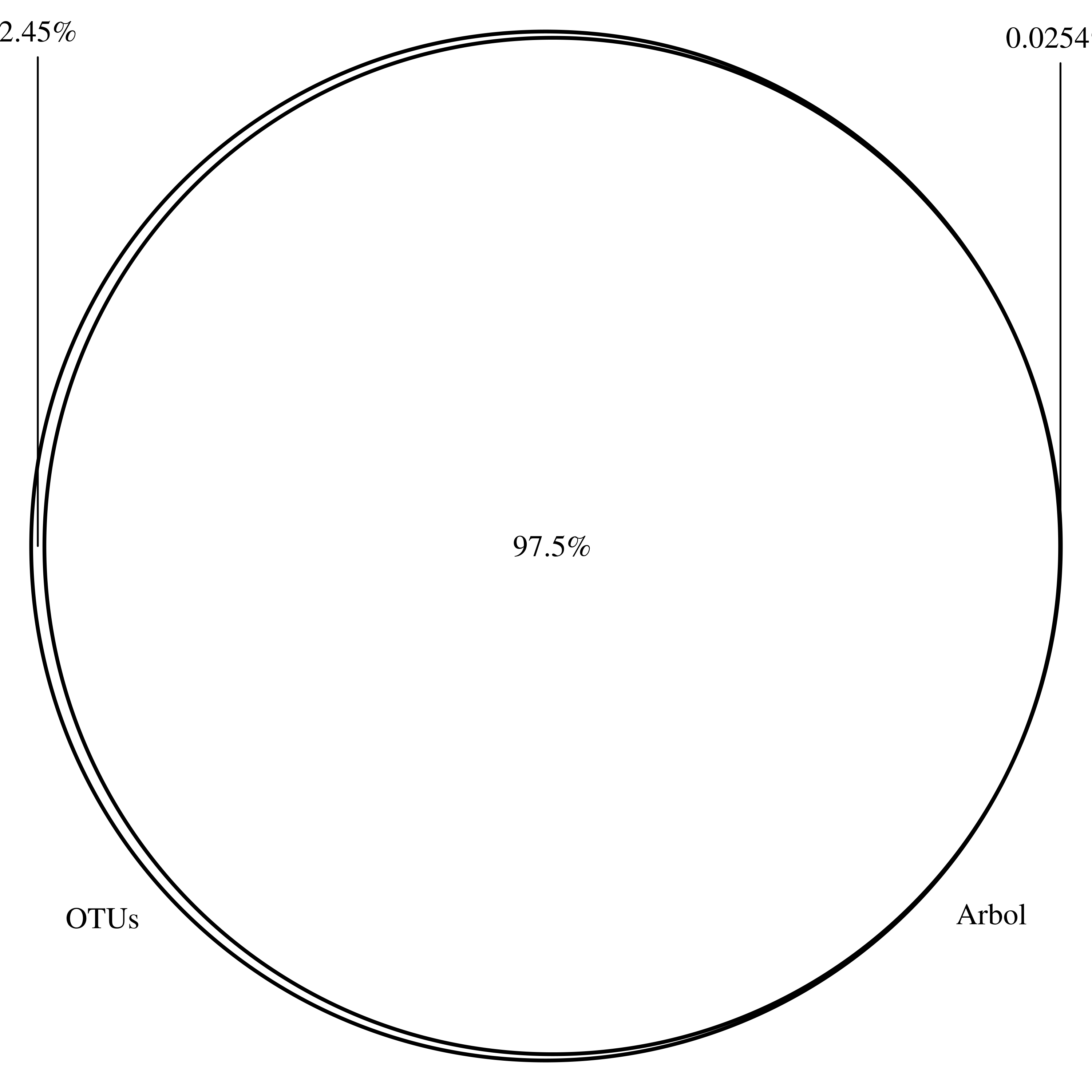

### Reads_cores_Tree_ALL_OTUs_reads_core_ALL_.png

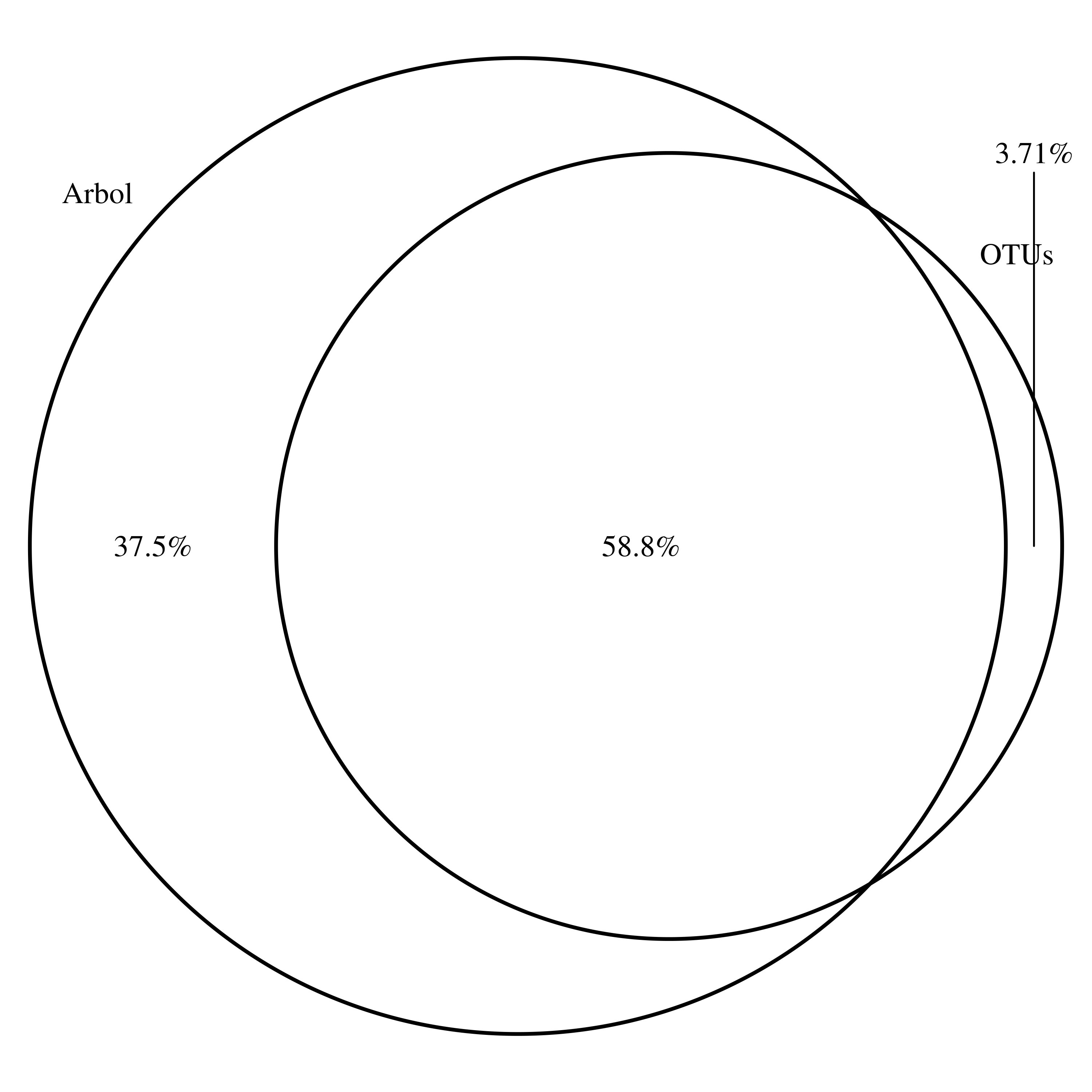

### Supplementary material 2

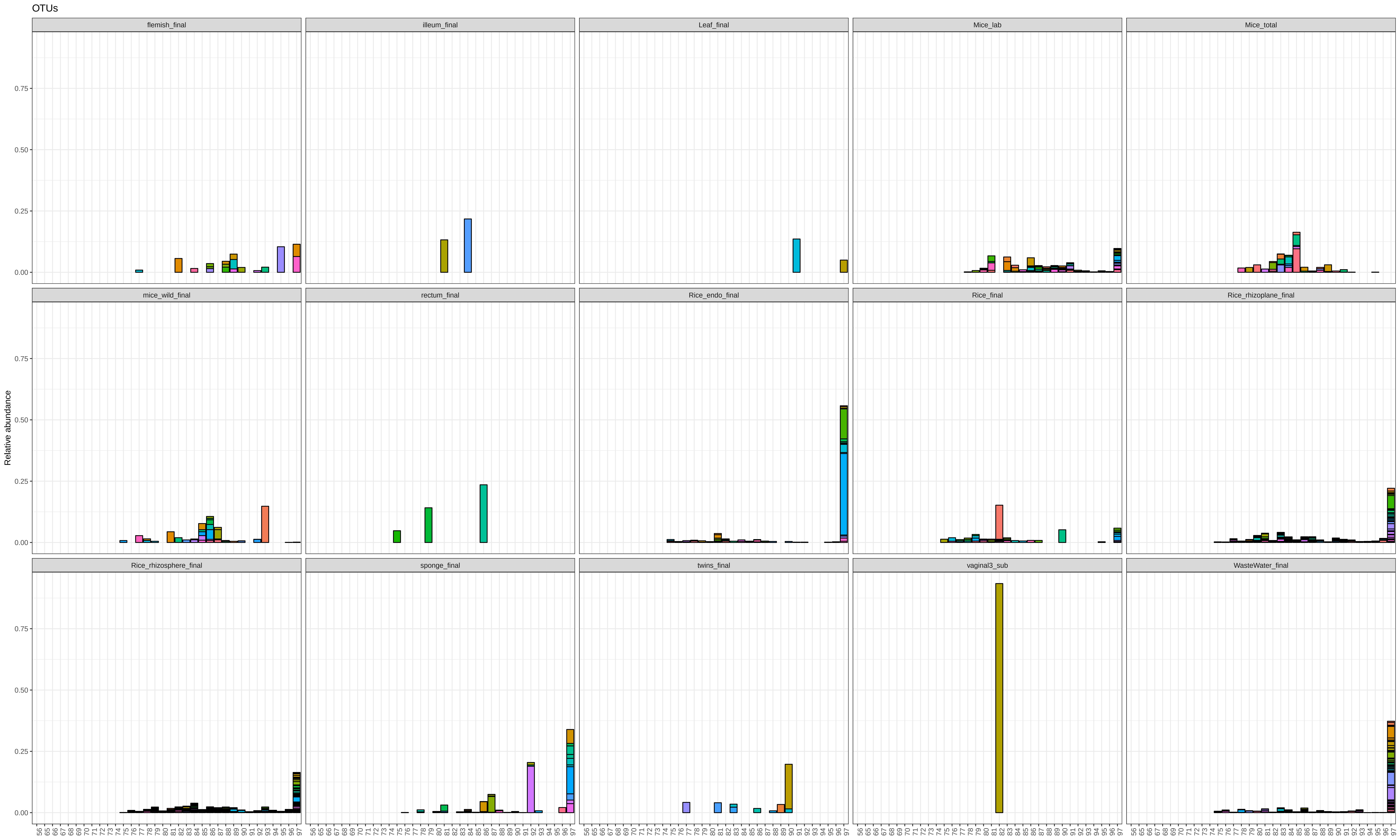

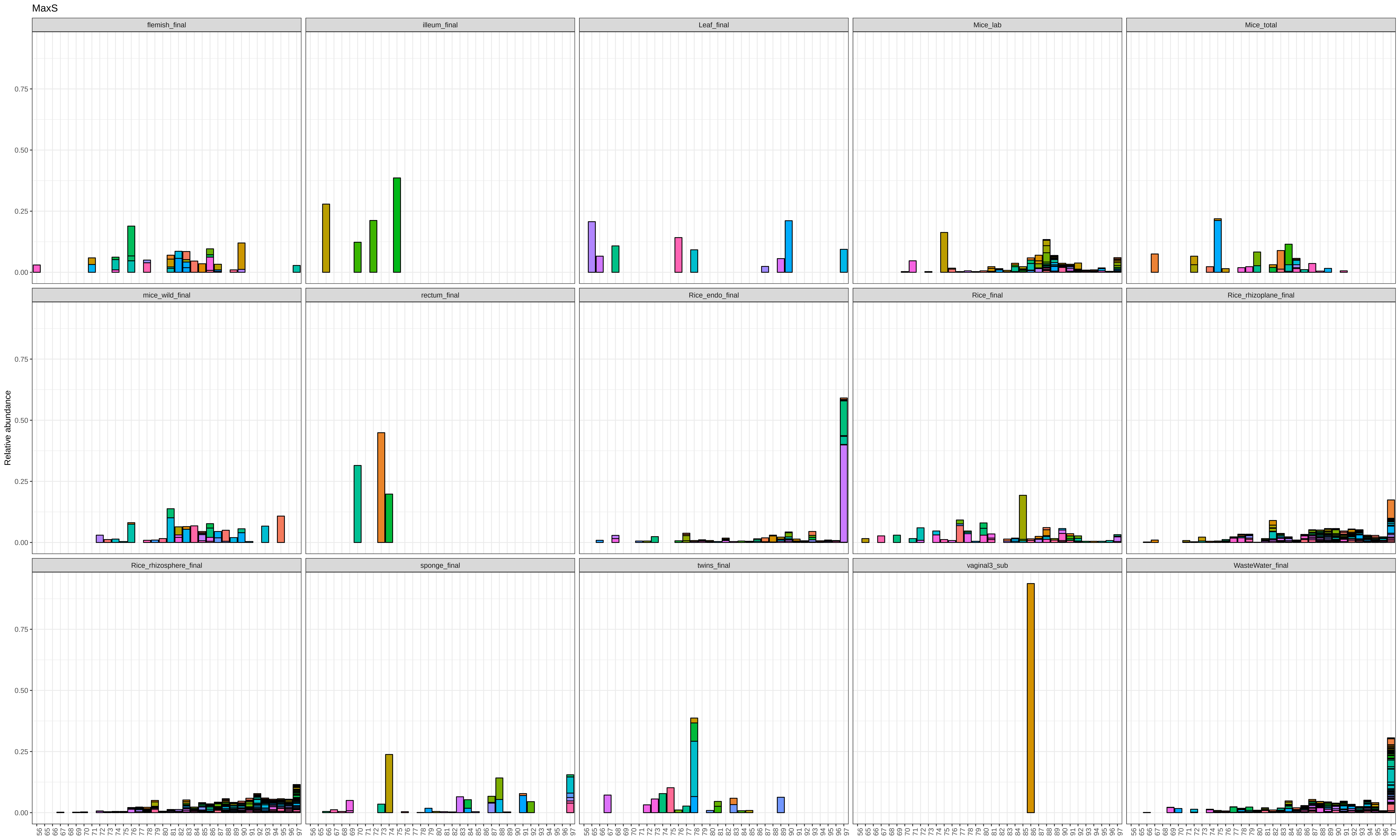

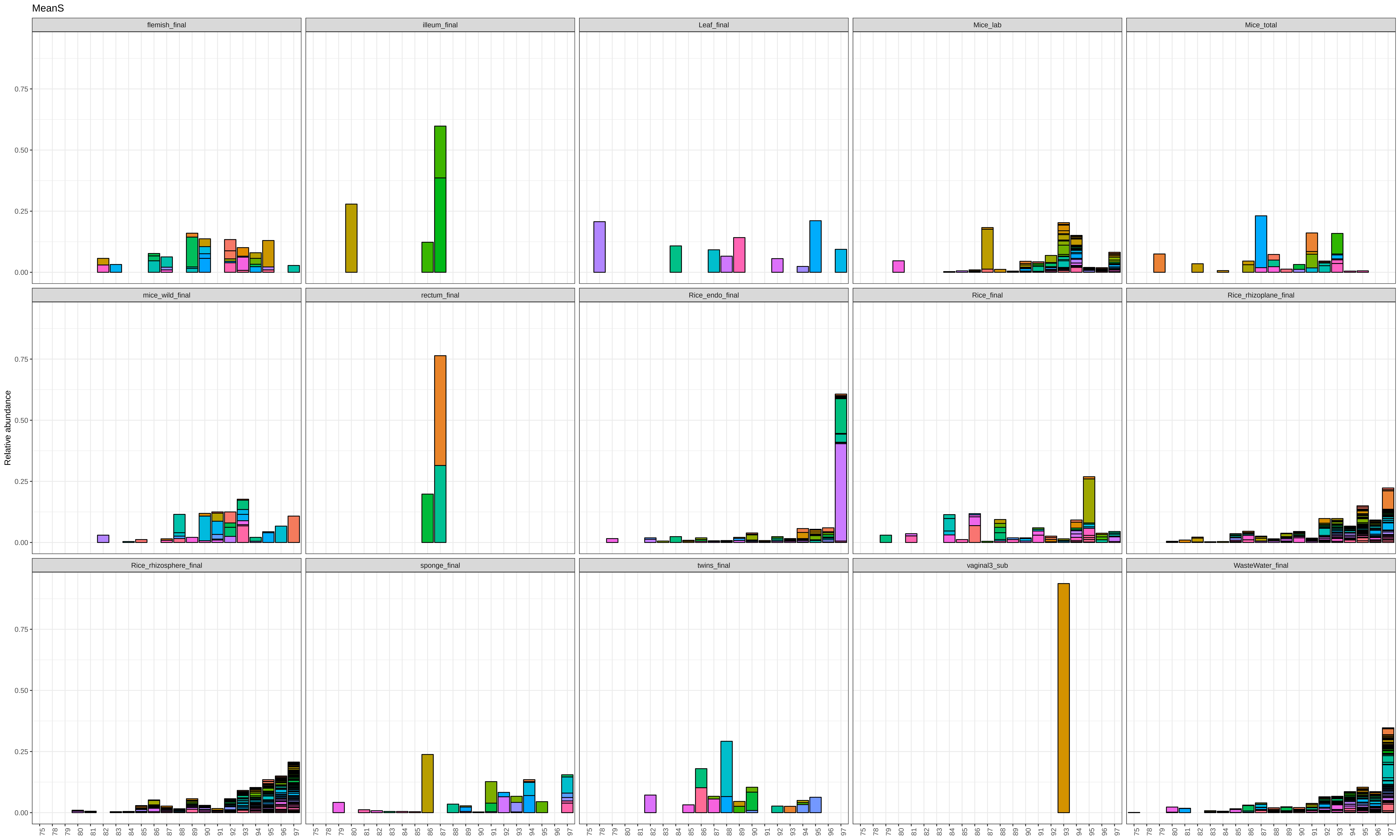

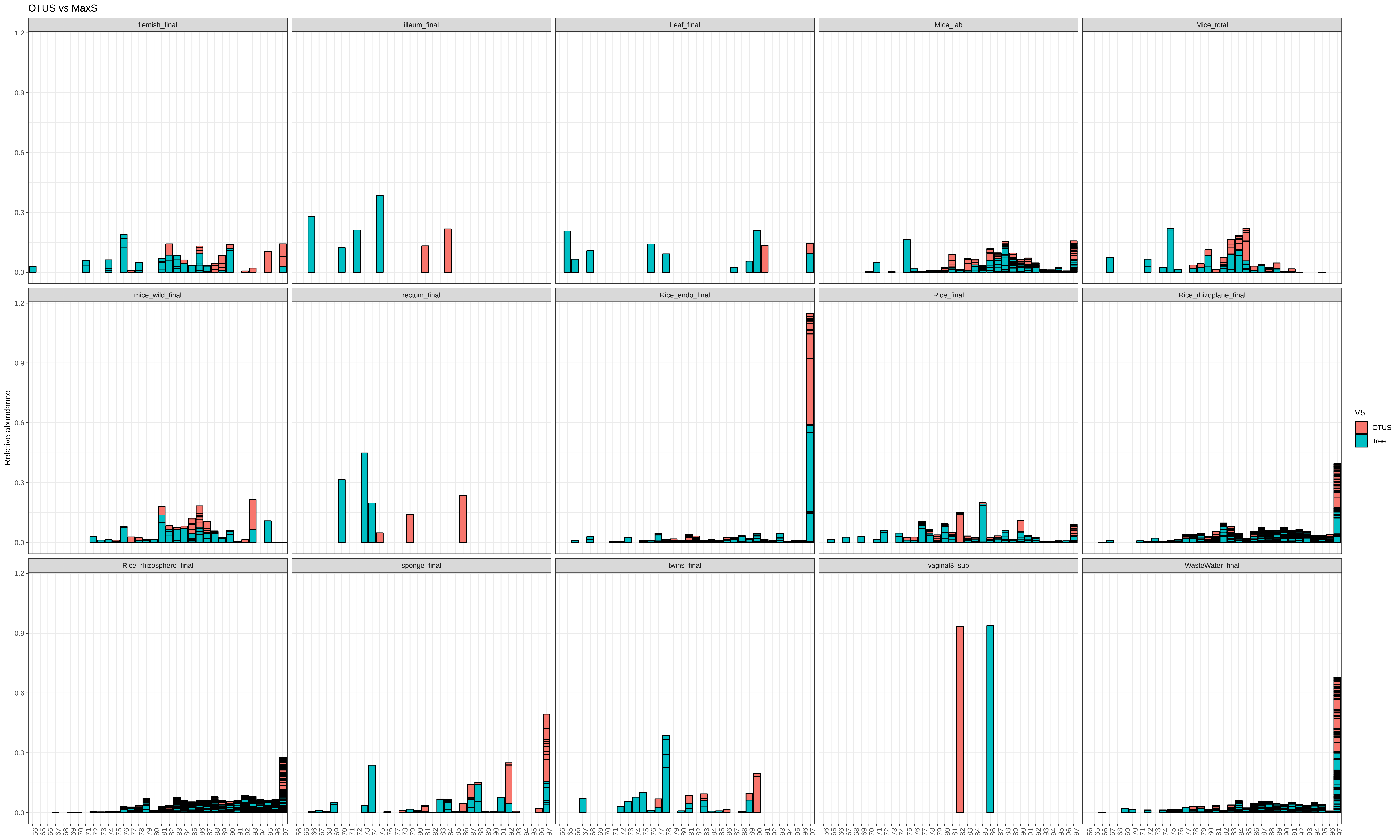

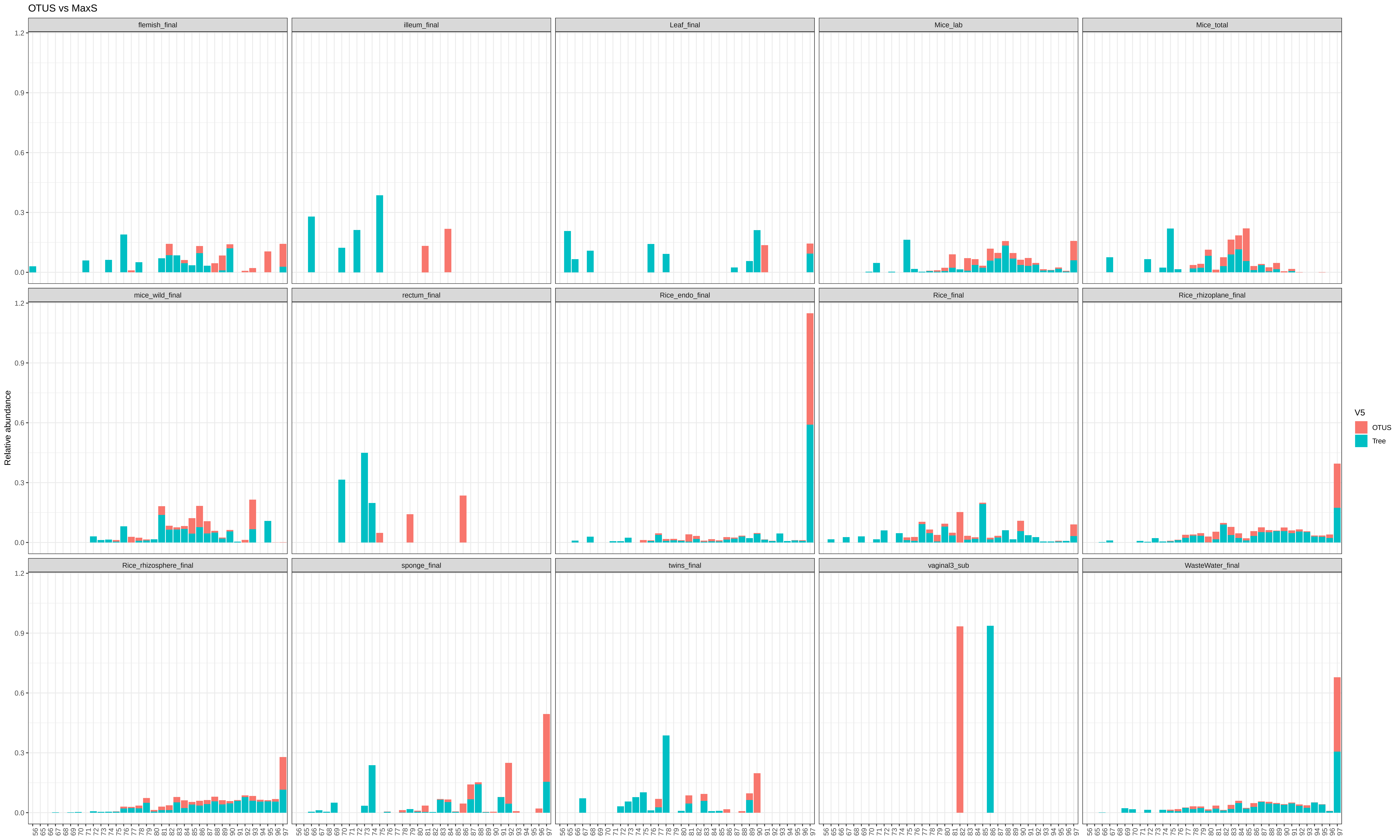
